# Construction of Animal Models Based on Exploring Pathological Features and Mechanisms of Different Locations in the Progression of DVT-APTE-CTEPD/CTEPH

**DOI:** 10.1101/2024.03.28.587300

**Authors:** Qinghuang Lin, Wenfeng Wang, Xiaoyun Chen, Jixiang Liu, Nan Shao, Qiuxia Wu, Xingyue Lai, Maohe Chen, Min Chen, Yijin Wu, Dawen Wu, Hongli Li, Peiran Yang, Yunxia Zhang, Zhu Zhang, Zhenguo Zhai, Chaosheng Deng

**Author notes:** Equal contributors (Qinghuang Lin, Wenfeng Wang, Xiaoyun Chen, Jixiang Liu). Correspondence to: Professor Chaosheng Deng, The First Affiliated Hospital of Fujian Medical University, 20 Chazhong Road, Fuzhou, Fujian 350004, People’s Republic of China.Professor Zhenguo Zhai,Department of Pulmonary and Critical Care Medicine, Center of Respiratory Medicine, China-Japan Friendship Hospital, Beijing, China. No. 2, East Yinghua Road, Chaoyang District, Beijing, China.100029. (Chaosheng Deng). (Zhenguo Zhai).

## Abstract

**Background:** Chronic thromboembolic pulmonary disease (CTEPD) and chronic thromboembolic pulmonary hypertension (CTEPH) are sequelae of acute pulmonary embolism (APE) and severely affect patients’ health and quality of life. The treatment of these conditions is challenging, and their underlying mechanisms remain unclear. The main reason for this is the lack of an animal model that can fully simulate the entire chain of DVT-APTE-CTEPD/CTEPH progression. The objective of this study is to construct an ideal animal model that simulates the major pathological changes of DVT-APTE-CTEPD/CTEPH and can be used for mechanistic exploration. We aim to compare the advantages and disadvantages of different modeling approaches and provide an experimental basis for investigating the mechanisms of pulmonary embolism chronicization at different stages of evolution.

*Methods and Materials:* We first evaluated the pathological changes in the pulmonary arterial intima stripping tissue of CTEPH patients. Animal models were established by multiple injections of thrombus columns through the internal jugular vein to simulate distal remodeling of the pulmonary artery. To simulate significant remodeling and fibrosis in the middle and distal segments of the pulmonary artery, thrombus columns were injected along with splenectomy. A CTEPD model with intimal fibrosis remodeling was successfully established by selectively injecting large thromboemboli into the pulmonary artery sites in large animals (dogs). A rat model with pathological manifestations of intimal fibrosis remodeling in the proximal end of the pulmonary artery was constructed using large thrombi combined with nitric oxide synthase inhibitors. An animal model of DVT was established using the inferior vena cava ligation method.

*Results:* According to the different pathological features and mechanisms observed in the progression of human DVT-APTE-CTEPD/CTEPH, we constructed animal models that conform to these pathological manifestations and mechanisms, each with its own advantages. Furthermore, the different methods used to construct animal models can be integrated and applied together.

*Conclusion:* Animal models constructed using different modeling methods can effectively simulate the pathological and physiological manifestations of the corresponding stages of chronic pulmonary embolism. Researchers can select the aforementioned models according to their specific research purposes, directions, and requirements.

## Introduction

Venous thromboembolism (VTE) includes deep venous thrombosis (DVT) and pulmonary thromboembolism (PTE), with most cases of acute pulmonary thromboembolism (APTE) resulting from the detachment of thrombi from DVT. After three months of standard anticoagulant therapy for PTE, 25%-50% of patients still have unresolved thrombi in the pulmonary arteries, leading to residual thrombus and organization. This process results in the remodeling of the pulmonary arteries, known as chronic thromboembolic pulmonary disease (CTEPD). When CTEPD is accompanied by resting pulmonary hypertension (PH) (i.e., PH > 20mmHg), it is referred to as chronic thromboembolic pulmonary hypertension (CTEPH). CTEPD and CTEPH significantly impact patients’ quality of life. While there is no statistical data on the development of CTEPD in patients with PTE, approximately 4% progress to CTEPH [1-3]. There is no consensus on the treatment of CTEPD without PH, but pulmonary artery endarterectomy (PEA) has shown good efficacy in some patients. PEA is currently the most effective treatment for CTEPH, but even after successful surgery, 25% of patients still experience residual pulmonary hypertension. Other treatments, such as pulmonary artery balloon angioplasty and targeted drugs, have shown certain efficacy for CTEPD/CTEPH, but repeated treatments can be expensive [4]. Observation of pulmonary artery endarterectomy specimens after PEA reveals complete organization and fibrotic changes in the pulmonary arterial intima, showing a "muscle-like" tissue with alternating red and yellow colors macroscopically. Closer to the lumen, there may still be small amounts of dark red thrombus-like tissue, indicating possible in-situ thrombus formation (as indicated by the white arrow in Figure 1A). Microscopically, organized thrombi and fresh thrombi coexist, and the pulmonary arterial intima is significantly thickened, with abundant deposition of extracellular matrix such as collagen and SMA+ myofibroblasts. In contrast, the distal vessels appear as complete yellow "fibrous-like" tissue, with predominant intimal remodeling (as indicated by the black arrow in Figure 1A), characterized by thickened pulmonary arterial intima, evident reticular and filamentous fibrosis. The region between the proximal and distal ends is referred to as the middle segment, which shows a combination of the characteristics observed in the proximal and distal segments. It exhibits intimal remodeling and fresh thrombus formation (as indicated by the blue arrow in Figure 1A).

**Figure 1A.**
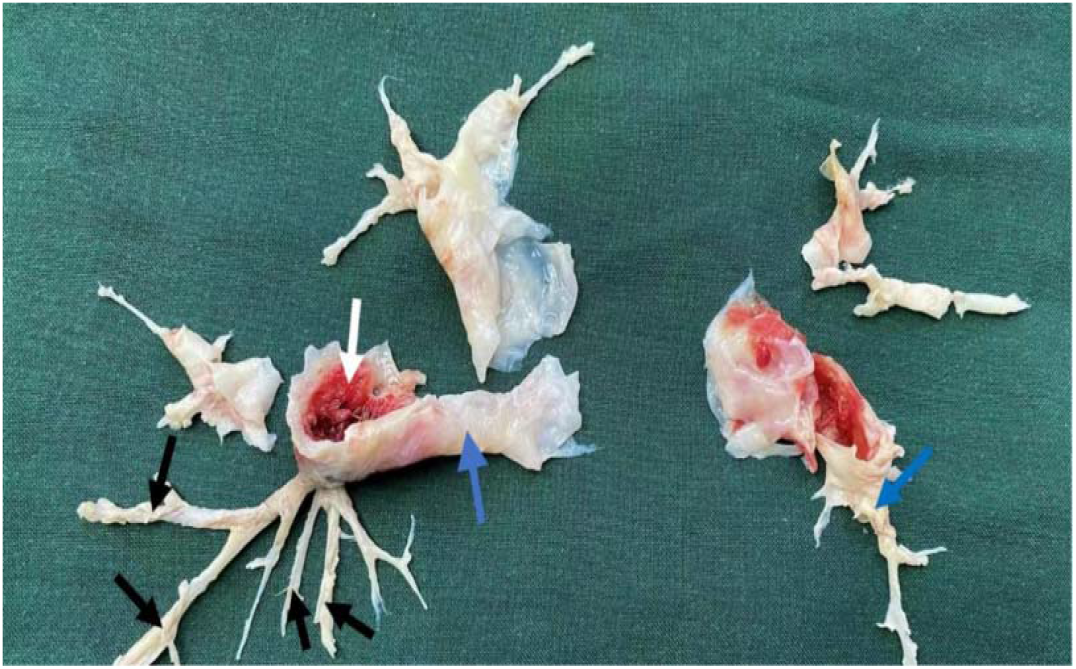
Histological Specimen of Pulmonary Arterial Intimal Remodeling in CTEPH Patients (Image from China-Japan Friendship Hospital, used with authorized permission) Note: The black arrows indicate thickening and remodeling of the distal pulmonary arterial intima. The blue arrows indicate significant vascular remodeling in the mid pulmonary arteries. The white arrows indicate the proximal pulmonary arterial intima, with a small amount of dark red thrombus-like tissue attached.

Currently, the treatment of CTEPD is challenging due to the lack of a clear understanding of its specific mechanisms of occurrence and progression. There is limited basic research on this topic, and previous studies have suggested that the key pathological changes in CTEPD are pulmonary artery thrombus fibrosis, followed by proximal pulmonary artery thrombus and intimal fibrosis, as well as middle and distal pulmonary artery intimal remodeling [5] (as shown in Figure 1B). However, there are few studies exploring the aforementioned pathological and physiological mechanisms of CTEPD, mainly due to the difficulty in establishing animal models for CTEPD. Previous attempts by researchers to establish animal models through repeated thrombus embolization have been unsuccessful [6,7], primarily due to the strong fibrinolytic capacity of animals, which leads to rapid dissolution of injected thrombi, preventing chronic remodeling. Some researchers have ligated the left pulmonary artery in piglets and repeatedly injected non-dissolvable microspheres through the external jugular vein to induce right pulmonary embolism and establish a CTEPD model. Although this model partially simulates the pathological and physiological characteristics of CTEPD [8], it fundamentally fails to completely simulate the mechanisms of pulmonary artery obstruction caused by the detachment of DVT thrombi and subsequent thrombus organization leading to pulmonary vascular remodeling.

**Figure 1B.**
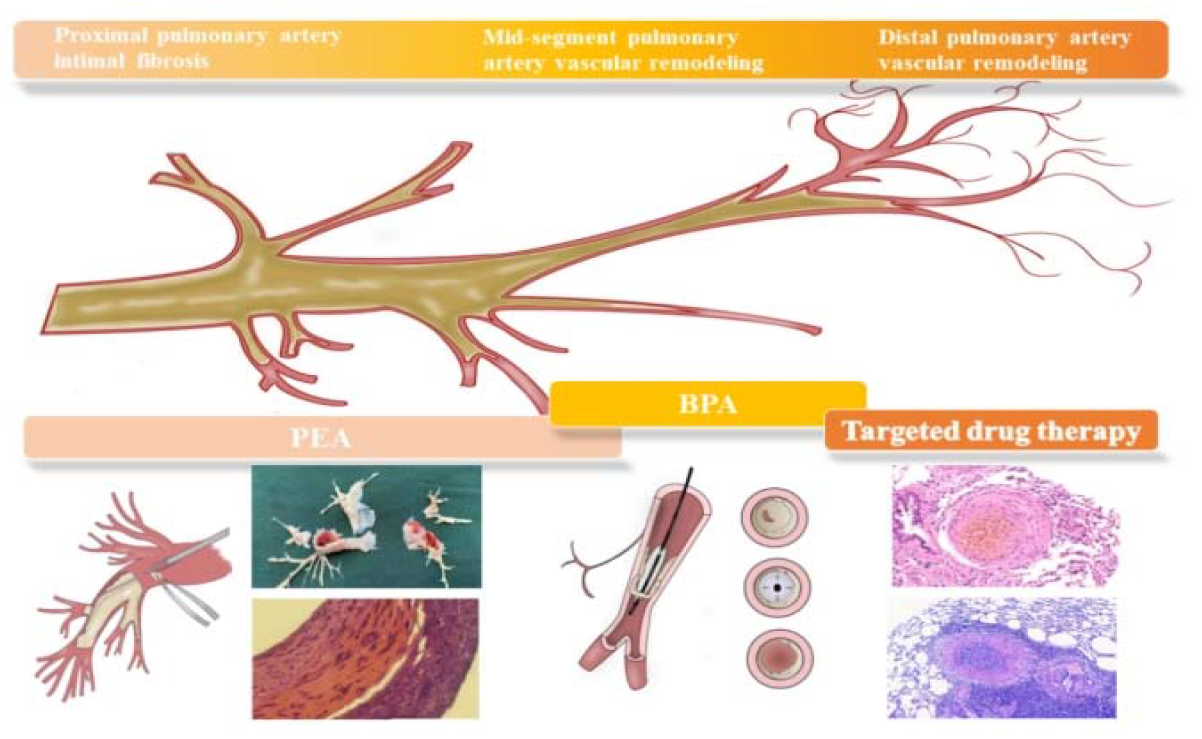
Pathological Changes in Chronic Pulmonary Embolism and Current Clinical Treatment Modalities.

Therefore, in this study, we provide a detailed description of the construction of animal models for the chronicization of pulmonary embolism with different pathological and physiological changes. During the process of constructing animal models for the chronicization of pulmonary embolism, we found that it is not feasible to fully simulate the pathological changes from the distal to proximal ends of the pulmonary arteries in a single animal due to the efficient fibrinolytic system. Therefore, we constructed animal models that simulate the specific pathological features and mechanisms of CTEPD at different locations along the pulmonary arteries, from the distal to proximal ends, observed in human CTEPD patients.

## Materials and Methods

### Human Pulmonary Artery Tissue Specimens

#### Control Group

Ten patients (4 males and 6 females) without pulmonary arterial hypertension or pulmonary embolism who underwent lung lobectomy surgery at the First Affiliated Hospital of Fujian Medical University were included in the control group. The age range was 48-70 years, with a mean age of 55 years. These patients were selected from individuals undergoing health check-ups. Inclusion criteria were as follows: (1) age between 30 and 75 years; (2) no evidence of pulmonary embolism or signs of pulmonary arterial hypertension based on examinations such as CT pulmonary angiography and color Doppler echocardiography. Exclusion criteria were as follows: (1) organ fibrotic diseases such as pulmonary fibrosis; (2) vasculitis diseases such as granulomatosis with polyangiitis; (3) unwillingness to cooperate.

### Chronicization of Pulmonary Embolism Group

Ten patients (3 males and 7 females) diagnosed with chronic thromboembolic pulmonary hypertension (CTEPH) according to the 2018 Chinese guidelines for the diagnosis, treatment, and prevention of pulmonary thromboembolism were included in this group. These patients underwent pulmonary endarterectomy surgery at Beijing Chaoyang Hospital, Capital Medical University, between April and December 2020. The age range was 48-70 years, with a mean age of 57 years. All patients underwent CT pulmonary angiography and were confirmed to have CTEPH with mean pulmonary artery pressure (mPAP) > 25 mmHg measured by right heart catheterization. The exclusion criteria were the same as those for the control group.

All the participants in the control and patient groups provided informed consent and received approval from the ethics committee.

### Construction of Chronic Pulmonary Embolism Animal Model

Overall plan: Establish an animal model that mimics the distal remodeling of the pulmonary artery by repeatedly injecting thrombus columns through the internal jugular vein. By injecting thrombus columns in combination with splenectomy, establish an animal model that simulates significant remodeling and fibrosis in the middle and distal segments of the pulmonary artery. The proximal endovascular fibrosis remodeling, which mimics human pulmonary endarterectomy (PEA) surgery, requires the use of large thrombi and multiple embolizations, which can easily lead to the death of experimental animals. Therefore, we established a chronic thromboembolic pulmonary hypertension (CTEPH) model in large animals, specifically dogs, by selectively injecting large thrombi into the pulmonary artery, resulting in residual thrombi and endovascular fibrosis remodeling. However, large animals such as dogs are expensive and require higher maintenance conditions. In order to facilitate the application of similar models, we also established a rat model using larger thrombi in combination with nitric oxide synthase inhibitors, which simulates the pathological manifestations of endovascular fibrosis remodeling in the proximal end of the pulmonary artery and increased pulmonary artery pressure. Furthermore, research has found that the biological effects of chronic pulmonary artery thrombosis are similar to deep vein thrombosis. Therefore, we further constructed an animal model of pulmonary embolism originating from deep vein thrombosis using the inferior vena cava ligation method, which simulates the thrombosis organization and recanalization of pulmonary embolism.

### Animal Source

Sixty male Sprague Dawley rats, aged 3 months, weighing 300-350g, and sixteen healthy adult mixed-breed dogs, of either sex, weighing (20.0±1.5) kg, were used. All rats and dogs were housed in animal rooms with a temperature of 20-24°C and humidity of 65-70%, with free access to food and water. All experimental and animal management protocols were conducted under the permission of the Animal Use and Management Committee of Fujian Medical University and the Chinese Academy of Medical Sciences, and in accordance with the guidelines for the use and care of laboratory animals in the United States.

### Construction of Mid and Distal Pulmonary Artery Remodeling Model

#### Experimental Animal Groups

Sixty Sprague Dawley rats were randomly divided into three groups: sham operation group (n=20), chronic pulmonary embolism group (n=20), and splenectomy + chronic pulmonary embolism group (n=20).

### Autologous Thrombus Preparation

After ether anesthesia, rats were used to collect blood via the retro-orbital vein using a capillary tube (inner diameter 1.0mm). The collected blood was placed in a sterile glass petri dish containing aminocaproic acid solution overnight. The next day, the solution in the dish was flushed out, and the thrombus was cut into 3mm-long columns using a sterile scalpel blade. The thrombus was drawn into a syringe and connected to a 7F syringe needle catheter for later use.

### Model Construction

#### Splenectomy in Rats

The hair below the left rib margin of rats was trimmed, and the skin was incised and separated layer by layer until the spleen was exposed. The spleen was removed, and the incision was sutured. The rats were allowed to recover naturally.

### Right External Jugular Vein Catheterization in Rats

External jugular vein catheterization was performed on rats with or without splenectomy after one week of feeding. The area around the neck of the rats was shaved. A longitudinal incision was made on the right side of the midline of the neck to expose the right external jugular vein, which was approximately 0.5cm in length. A 3.0 silk suture was used to loosely tie a knot at the proximal end of the right external jugular vein, and the distal end of the vein was ligated. A "V"-shaped incision was made with an ophthalmic scissors on the distended segment of the jugular vein. The catheter was inserted into the external jugular vein through this incision, approximately 2cm deep, and the catheter was secured with a ligature looped around the fixed knot. Blood was withdrawn and injected with physiological saline to check for successful catheter placement, indicated by no resistance or leakage. The rat was placed in a prone position, and two small incisions of 0.5cm were made behind both ears. A glass spatula was used to create a subcutaneous tunnel next to the right external jugular vein, which emerged from the small incision behind the neck. The distal end of the catheter was grasped with forceps and pulled out along the subcutaneous tunnel. A 3.0 silk suture was used to tie a knot around the catheter, securing it in the subcutaneous tissue, with 2cm of the catheter left outside. The indwelling needle tubing was placed into the catheter under the guidance of the needle core, and the needle core was removed and connected to a heparin cap. The indwelling needle catheter was placed subcutaneously, with the heparin cap outside, and the exit of the catheter was sutured for fixation.

### Thrombus Injection

The day after catheter placement, blood was withdrawn from the catheter using a syringe to confirm blood flow. Autologous blood clots prepared in advance were injected into the experimental group rats through the right external jugular vein catheter, with continuous administration of thrombus and intramuscular injection of aminocaproic acid throughout the process. After two weeks, the rats underwent thoracotomy, the left lung tissue was collected, and the pulmonary arteries with thrombotic occlusion were isolated.

### Construction of Pulmonary Arterial Endothelial Fibrosis Model

#### Construction of Pulmonary Arterial Endothelial Fibrosis Model in Rats

##### Experimental Animal Groups

Sixty SD rats were randomly divided into three groups: sham operation group (n=20), NG-Nitro-L-arginine Methyl Ester (L-NAME) control group (n=20), and chronic pulmonary embolism group (n=20). The chronic pulmonary embolism group received intraperitoneal injections of 50 mg/kg·d L-NAME for one week prior to thrombus injection, until the day of pulmonary artery tissue collection. On the eighth day of L-NAME administration, 5 autologous blood clots measuring 3 mm in length and 1.3 mm in diameter, suspended in 2 mL of aminocaproic acid solution, were injected into the rats through the left external jugular vein using a 9F needle connected to a syringe. Repeat injections were performed 4 and 7 days later. In the sham operation group, an equal volume of normal saline was injected instead of autologous blood clots. In the L-NAME group, in addition to normal saline replacing the thrombus injection, L-NAME was intraperitoneally injected daily.

### Measurement of Pulmonary Arterial Pressure and Specimen Collection

The right external jugular vein of rats was isolated, and a PE-50 polyvinyl chloride catheter connected to a multi-channel physiological recorder was inserted. The pressure curve was observed, and the position of the catheter was adjusted until it was inserted into the pulmonary artery. Pulmonary arterial pressure was measured and recorded. After pressure measurement, thoracotomy was performed, and the left lung tissue was collected. The pulmonary arteries with thrombotic occlusion were identified. Based on the location of thrombotic occlusion in the chronic pulmonary embolism rat group, lung tissue from the sham operation group and L-NAME group at the same level was collected.

### Construction of Canine Pulmonary Arterial Endothelial Fibrosis Model

#### Experimental Animal Grouping

Sixteen healthy adult mixed-breed dogs were used. Among them, 15 dogs with occlusion of the left lower pulmonary artery were divided into three groups, with 5 dogs in each group: sham operation group; 1-week group with occlusion of 5 thrombus columns observed for 1 week; 2-week group with occlusion of 5 thrombus columns observed for 2 weeks. In addition, one dog had thrombus column occlusion in the right lower pulmonary artery to confirm the feasibility of selective occlusion, and it was observed for 2 weeks.

### Establishment of Canine Pulmonary Arterial Endothelial Fibrosis Animal Model

Autologous venous blood from the dogs was used to create three thrombi: one approximately 7 cm long and 4.5 mm in diameter, and two approximately 5 cm long and 5.5 mm in diameter. These thrombi were aspirated into a long plastic catheter (Catheter I) for later use. The experimental dogs were intravenously injected with 5 ml of 1% propofol and then intraperitoneally injected with 3% pentobarbital sodium at a dose of 0.5 ml/kg for anesthesia. Endotracheal intubation was performed. A 7F sheath Swan-Ganz floating catheter (from Arow, USA) with a length of 50 cm and an inner diameter of approximately 5.5 mm (Catheter II) was prepared. The right external jugular vein was incised, and a 7F sheath was inserted. The Swan-Ganz floating catheter was inserted through the sheath. The floating catheter was placed inside Catheter II, and under X-ray guidance, Catheter II was selectively positioned at the common trunk of the middle lobe and the lower lobe. Thrombus columns in Catheter I were quickly injected into the corresponding pulmonary artery through Catheter II. Throughout the process, 0.3 g of aminocaproic acid was administered via intramuscular injection twice a day.

### Pathological Observation of Occluded Lung Tissue and Thrombus

At 1 and 2 weeks after modeling, the animals were euthanized using rapid exsanguination. The pulmonary artery was longitudinally opened along its course, and lung tissue containing thrombi was collected.

Construction of Rat Deep Vein Thrombosis (DVT) Model by Inferior Vena Cava Ligation Twenty SD rats were used. They were fasted before surgery, and a midline incision was made to access the abdomen. The inferior vena cava and its branches were isolated, and the inferior vena cava below the left renal vein was ligated using 4-0 silk sutures. The layers of tissue were sutured, and the rats were allowed to recover naturally. After 1 week, thrombus tissue in the rat inferior vena cava was collected.

### Pathological Tissue Examination

#### Histopathology

Tissues from patients with chronic thromboembolic pulmonary hypertension (CTEPH), control pulmonary arteries, rat pulmonary arteries, canine pulmonary arteries, and rat inferior vena cava thrombi were fixed in 4% paraformaldehyde for 48 hours. After dehydration and paraffin embedding, sections were stained with Hematoxylin and Eosin (HE), Sirius Red, Phosphotungstic Acid Hematoxylin (PTAH), and Masson’s trichrome. The pathological changes in the tissues were observed using an optical microscope (Leica DMI3000M, Germany).

### Immunofluorescence

Sections of human pulmonary artery tissue and rat pulmonary artery tissue were deparaffinized and rehydrated, followed by incubation in 0.3% Triton X-100. PBS wash was performed, and then gastric protease was added and incubated at 37°C for 20 minutes. PBS wash was performed again for decolorization. A 3% H2O2 solution was added to the slides and incubated at room temperature for 25 minutes, followed by washing with PBS. Then, 3% BSA was added and incubated at 37°C for 30 minutes for blocking. The primary antibodies CD45 (1:100) and ColL (1:150) were added to the slides and incubated overnight at 4°C. PBS wash was performed, and the secondary antibodies were added. The slides were incubated at 37°C in the dark for 30 minutes, followed by PBS wash, air-drying, and mounting with an anti-fluorescence quenching mounting medium.

### Immunohistochemistry

Sections of human pulmonary artery tissue and rat pulmonary artery tissue were deparaffinized and rehydrated. After blocking and antigen retrieval, the sections of human pulmonary artery tissue were respectively treated with SAP (1:150), IL-4 (1:100), IL-13 (1:200), TGF-β1 (1:300), Elastin (1:200), ColL (1:100), and α-SMA (1:200). The sections of rat pulmonary artery tissue were respectively treated with Elastin (1:200) and α-SMA (1:200). The slides were incubated overnight at 4°C. After rinsing, DAB staining, nuclear counterstaining with Hematoxylin, dehydration, clearing, and mounting were performed.

### Measurement of rats plasma NO levels

NO levels in plasma of chronic pulmonary embolism groups and controls were measured using specific ELISA according to the manufacturer’s instructions(Aimeng Youning, Shanghai).

### Proteomic Analysis of Human Pulmonary Artery Tissue

#### Protein Extraction

Take an appropriate amount of human pulmonary artery tissue sample and grind it into powder in a mortar with liquid nitrogen. Add four times the volume of the powder of lysis buffer (8M urea, 1% protease inhibitor, and 2 mM EDTA) to each sample and sonicate for lysis. After centrifugation, transfer the supernatant to another centrifuge tube and measure the protein concentration using a BCA assay kit.

#### Enzymatic Digestion

Add dithiothreitol (DTT) to the protein solution to a final concentration of 5 mM and reduce at 56°C for 30 minutes. Then add iodoacetamide to a final concentration of 11 mM and incubate at room temperature in the dark for 15 minutes. Finally, dilute the urea concentration in the sample to below 2M. Add trypsin at a mass ratio of 1:50 (trypsin:protein) and incubate overnight at 37°C. Continue the digestion by adding trypsin at a mass ratio of 1:100 (trypsin:protein) and incubate for an additional 4 hours.

#### TMT Labeling

The trypsin-digested peptides are dried by vacuum freeze-drying with Strata X C18 (Phenomenex). Dissolve the peptides in 0.5 M TEAB and thaw the labeling reagent, dissolve it in acetonitrile, and mix with the peptides. Incubate at room temperature for 2 hours and then vacuum freeze-dry.

#### HPLC Fractionation

The peptides are fractionated using an HPLC gradient of 8%-32% acetonitrile at pH 9 over a 60-minute period, resulting in 60 fractions. Then, the fractions are combined into 9 fractions, dried by vacuum freeze-drying, and subjected to further analysis.

#### Liquid Chromatography-Tandem Mass Spectrometry (LC-MS/MS) Analysis

The peptides are dissolved in mobile phase A (0.1% (v/v) formic acid in water) and separated using an EASY-nLC 1000 ultra-high-performance liquid chromatography system. Mobile phase A is 0.1% formic acid and 2% acetonitrile in water, while mobile phase B is 0.1% formic acid and 90% acetonitrile in water. The liquid phase gradient is set as follows: 0-65 min, 5%-16% B; 65-85 min, 16%-26% B; 85-87 min, 26%-70% B; 87-90 min, 70% B, with a flow rate of 300 nL/min. After separation by the ultra-high-performance liquid chromatography system, the peptides are ionized in the NSI ion source and then analyzed by Orbitrap Fusion Lumos mass spectrometry. Data acquisition is performed using a data-dependent acquisition (DDA) method.

#### Database Search

The second-level mass spectrometry data is searched using Maxquant (v1.5.2.8). The search parameters are set as follows: the database is SwissProt Human (20317 sequences), and a decoy database is added to calculate the false discovery rate (FDR) caused by random matches. The common contaminant database is also included in the search to eliminate the influence of contaminating proteins in the identification results. The enzyme specificity is set to Trypsin/P, allowing for up to 2 missed cleavage sites. The minimum peptide length is set to 7 amino acid residues, and the maximum number of modifications per peptide is set to 5. The mass tolerance for the first search and main search precursor ions is set to 20 ppm and 5 ppm, respectively, while the mass tolerance for fragment ions is set to 0.02 Da. Cysteine carbamidomethylation is set as a fixed modification, and variable modifications include methionine oxidation and protein N-terminal acetylation. The quantification method is set to TMT-10plex, and the FDR for protein identification and peptide-spectrum match (PSM) identification is set to 1%.

#### Statistical Analysis

Statistical analysis is performed using SPSS 25.0 software. Continuous variables are presented as (x ± s). One-way analysis of variance (ANOVA) is used for comparisons among multiple groups. If the assumption of equal variances is met, post hoc comparisons are performed using the Least Significant Difference (LSD) method. If the assumption of equal variances is violated, the Games-Howell method is used for comparisons. A P-value less than 0.05 is considered statistically significant.

## Results

### Clinical Specimen Study Results

#### Pathological and Fibrosis Examination

CTEPH patients showed significant pathological changes in the pulmonary artery intima, characterized by intimal thickening and proliferation of spindle-shaped fibroblasts (Figure 2). Sirius red staining and immunohistochemical staining for α-SMA, Collagen I, and Elastin demonstrated the deposition of extracellular matrix components, such as collagen fibers, in the pulmonary artery intima of CTEPH patients, indicating significant fibrosis (Figure 3).

**Figure 2.**
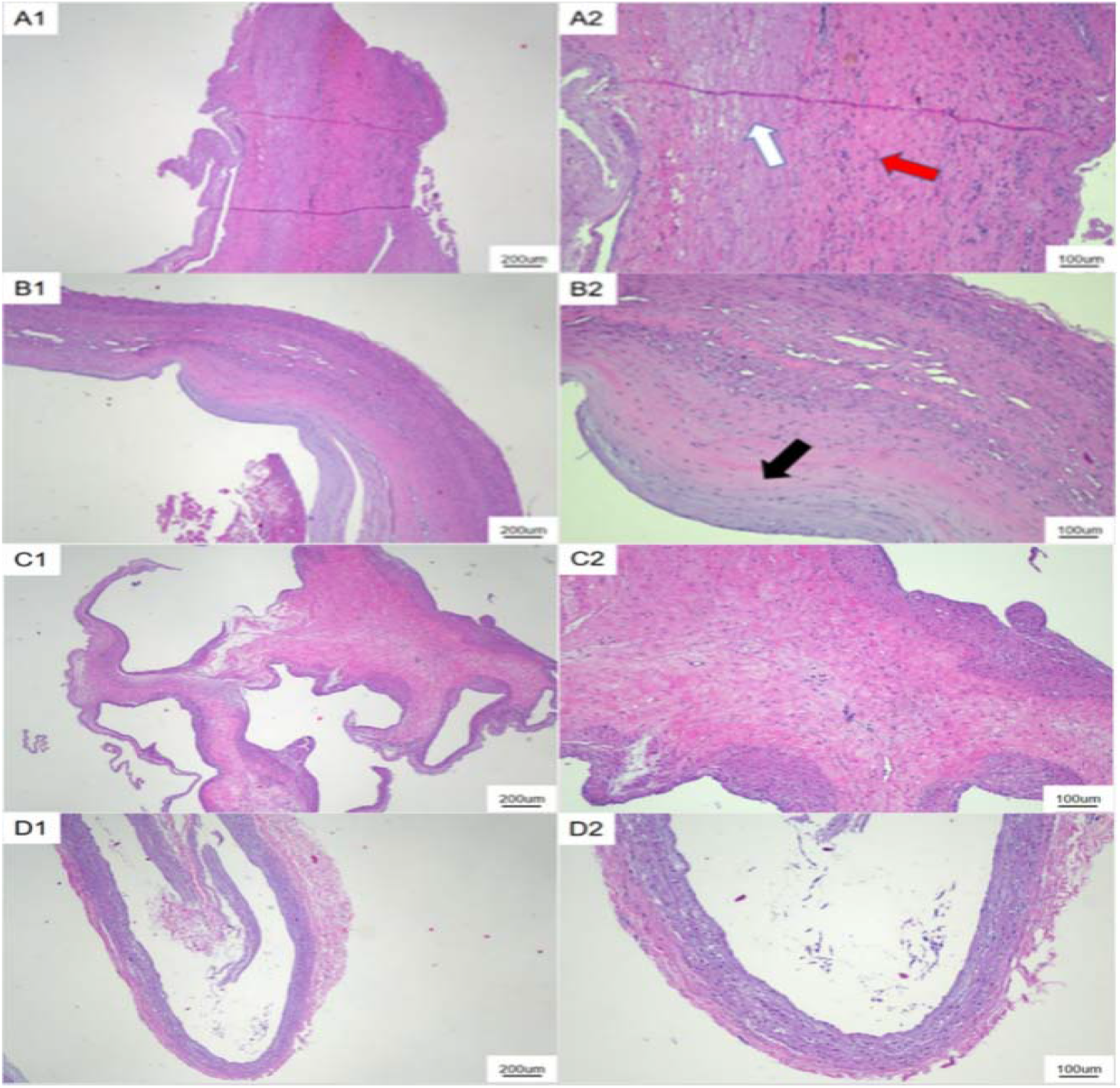
HE Staining Results of Human Pulmonary Artery Tissue. Note: A-D: HE staining results of human pulmonary artery tissue. A-C: HE staining results of pulmonary arterial intimal stripping tissue in CTEPH patients, where magnification factor 1 is 40x and magnification factor 2 is 100x for all serial numbers. D: HE staining results of control group pulmonary artery tissue, where magnification factor 1 is 40x and magnification factor 2 is 100x for all serial numbers.

**Figure 3.**
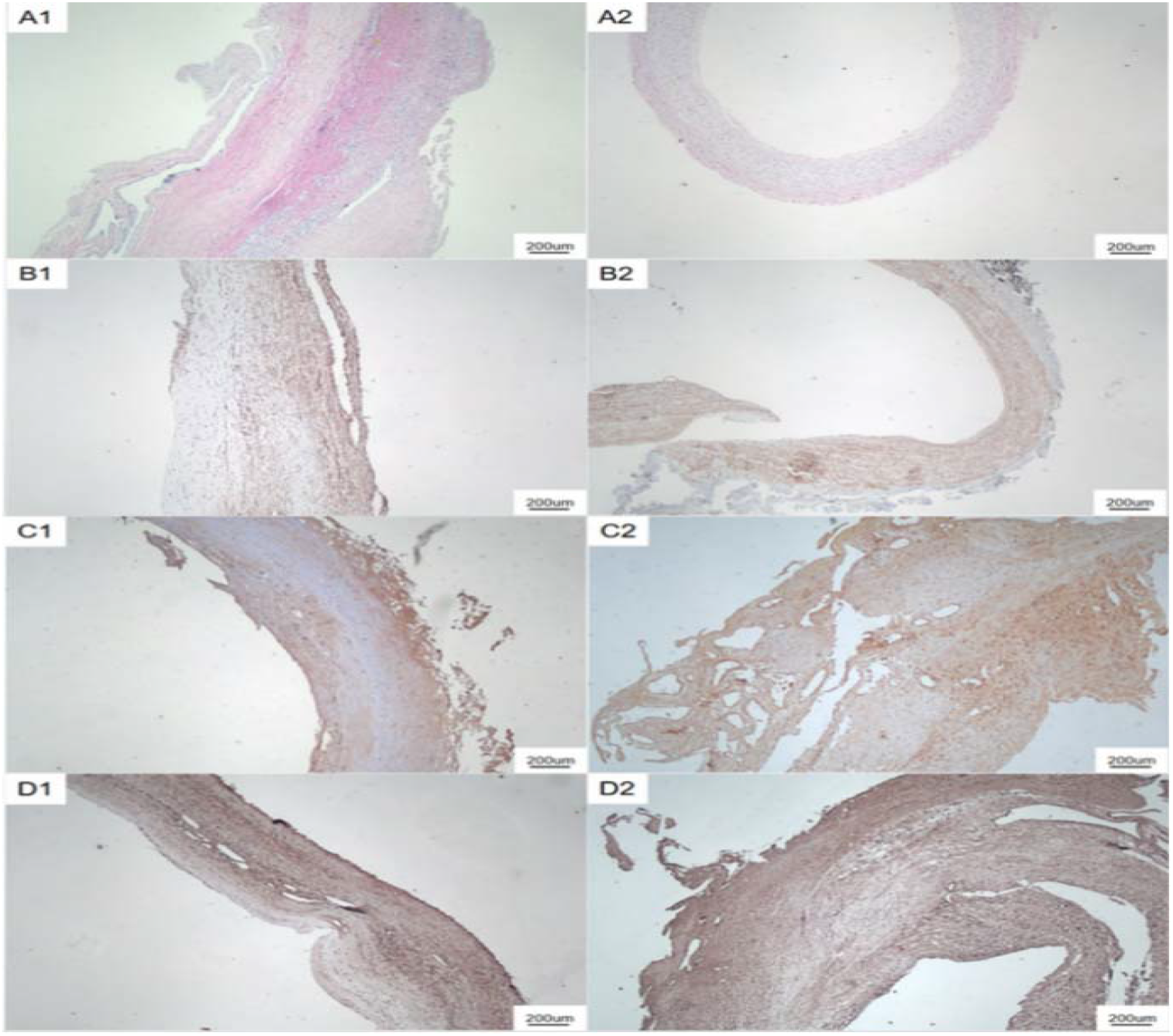
Extracellular Matrix Detection in Human Pulmonary Artery Tissue. Note: A1: Sirius Red staining of pulmonary arterial intimal stripping tissue in CTEPH patients (40x); A2: Sirius Red staining of control group pulmonary artery tissue (40x); B1: α-SMA immunohistochemical staining of pulmonary arterial intimal stripping tissue in CTEPH patients (40x); B2: α-SMA staining of control group pulmonary artery tissue (40x); C1-C2: Collagen I immunohistochemical staining of pulmonary arterial intimal stripping tissue in CTEPH patients (40x); D1-D2: Elastin immunohistochemical staining of pulmonary arterial intimal stripping tissue in CTEPH patients (40x).

### Proteomic Study

We performed high-throughput proteomic analysis on pulmonary artery tissues from 8 CTEPH patients and 8 control group patients. The results showed elevated expression of extracellular matrix proteins, such as Collagen α-1 (XII) chain and Collagen α-1 (V) chain, as well as elastin, in the pulmonary artery intima of CTEPH patients. The expression of anti-fibrotic protein SAP was decreased, while the expression of pro-fibrotic protein TGF-β1 was upregulated (Figure 4, Table 1). Data from Figure 5 and Figure 6 indicated that our proteomic experiments met quality control requirements and the data were reliable.

**Figure 4.**
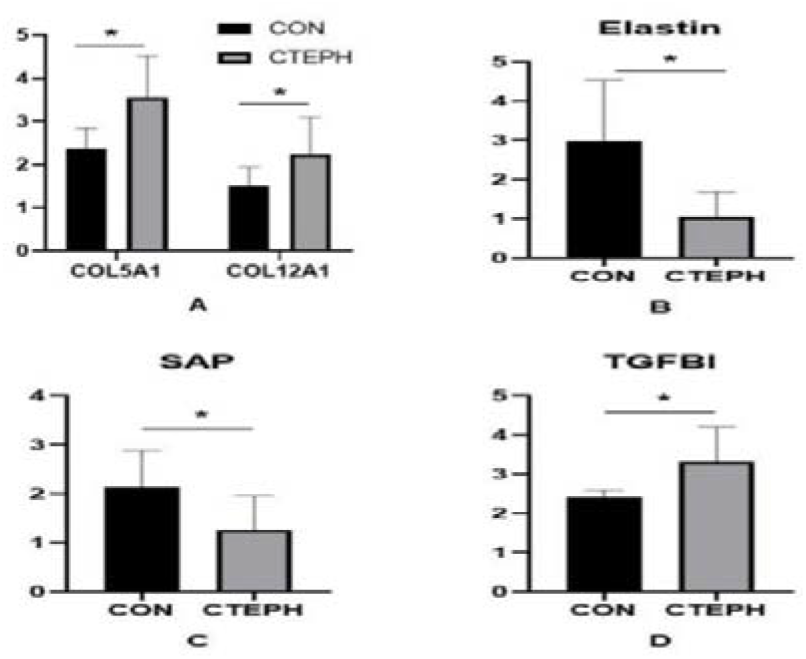
Expression of Extracellular Matrix and Fibrosis-Related Factors. Note: CON: Control group; CTEPH: Chronic Thromboembolic Pulmonary Hypertension group; Collagen alpha-1(XII) chain; Collagen alpha-1(V) chain; Elastin; SAP: Serum Amyloid P; Transforming growth factor-beta-induced protein ig-h3. *P<0.05.

**Figure 5.**
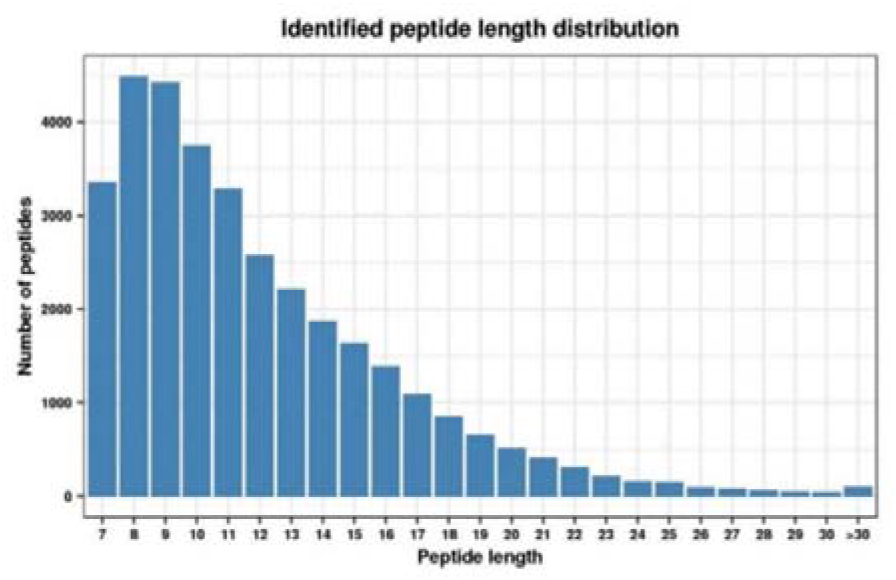
Length Distribution of Peptide Fragments Identified by Mass Spectrometry.

**Figure 6.**
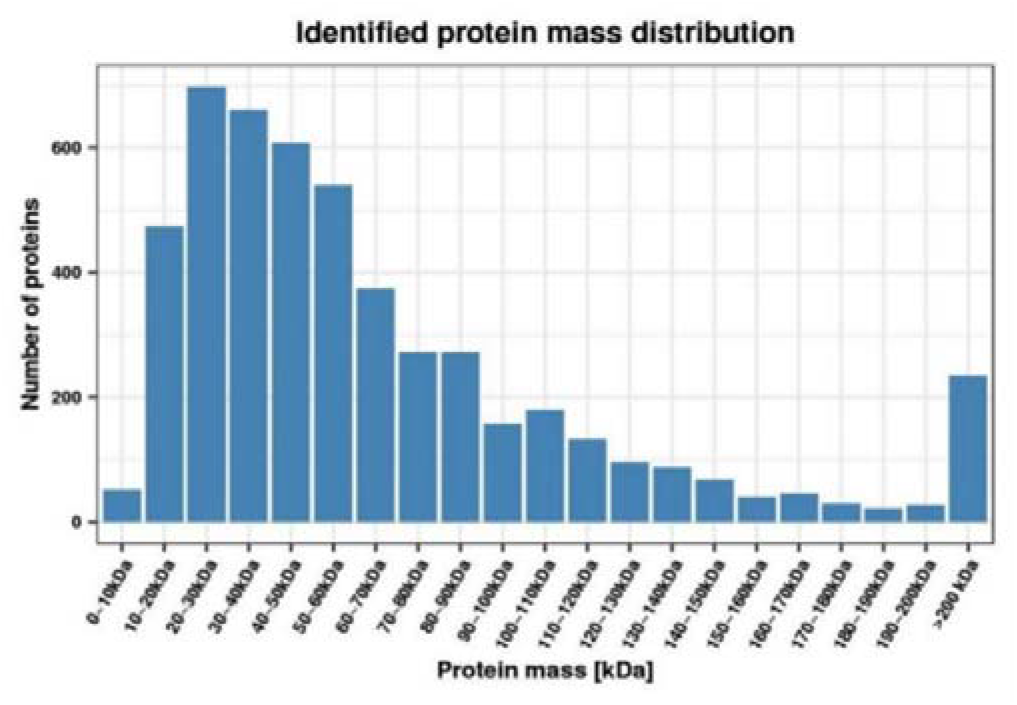
Molecular Weight Distribution of All Identified Proteins.

**Table 1.**
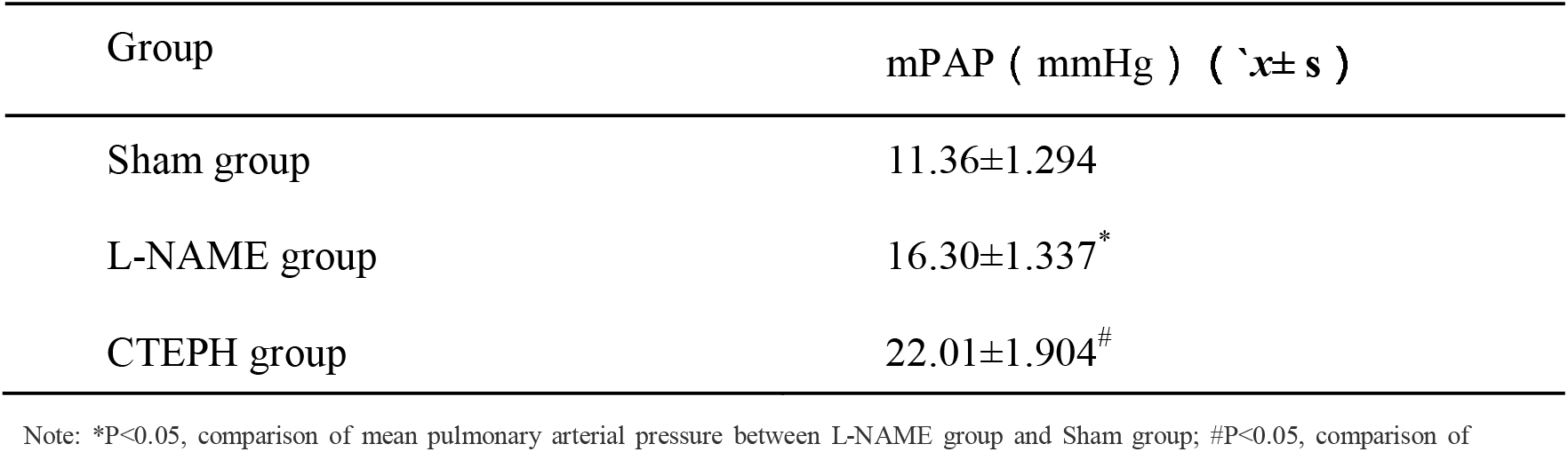
Expression Profile of Extracellular Matrix and Fibrosis-Related Factors.

### Immunofluorescence and Immunohistochemistry

Immunofluorescence results showed the presence of a large number of circulating fibroblasts co-expressing CD45 and Collagen I in the pulmonary artery intima of CTEPH patients (Figure 7). This preliminary finding suggests the involvement of circulating fibroblasts in pulmonary artery intimal fibrosis in CTEPH. We performed immunohistochemical staining using IL-4, IL-13, SAP, and TGF-β1 antibodies on pulmonary artery tissues from CTEPH patients (n=10) and the control group (n=10). The expression of IL-13 in the stripped pulmonary artery intima of CTEPH patients was significantly increased compared to the control group (Figure 8), indicating that IL-13 may be a key factor in the transformation of monocytes into circulating fibroblasts in the pulmonary artery intima.

**Figure 7.**
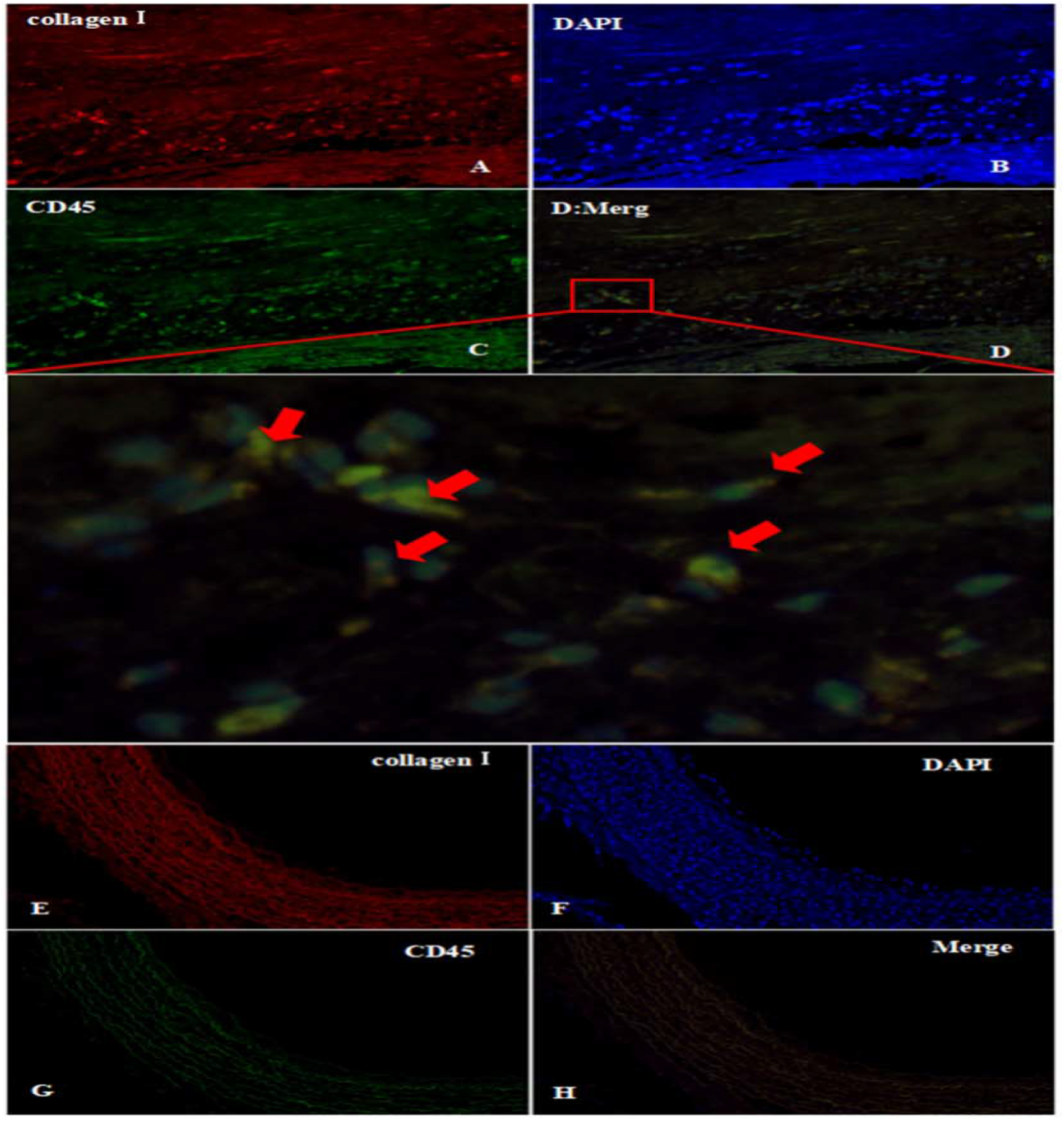
Immunofluorescence Double Staining of CD45 and Collagen I in Human Pulmonary Artery Intimal Tissue. Note: A-D: Immunofluorescence double staining of pulmonary arterial intimal stripping tissue in CTEPH patients. A: Collagen I (red); B: DAPI (4’,6-diamidino-2-phenylindole, a fluorescent dye that binds strongly to DNA) (blue), showing cell nuclei; C: CD45 (green); D: Merge; magnification factor is 200x for all. As indicated by the red arrow in panel D, cells positive for both CD45 and Collagen I can be observed, consistent with the aforementioned FC (fluorescence colocalization). E-H: Immunofluorescence double staining of control group pulmonary artery tissue. E: Collagen I (red); F: DAPI (blue); G: CD45 (green); H: Merge; magnification factor is 100x for all. No cells psitive for both CD45 and Collagen I are observed in panel H.

**Figure 8.**
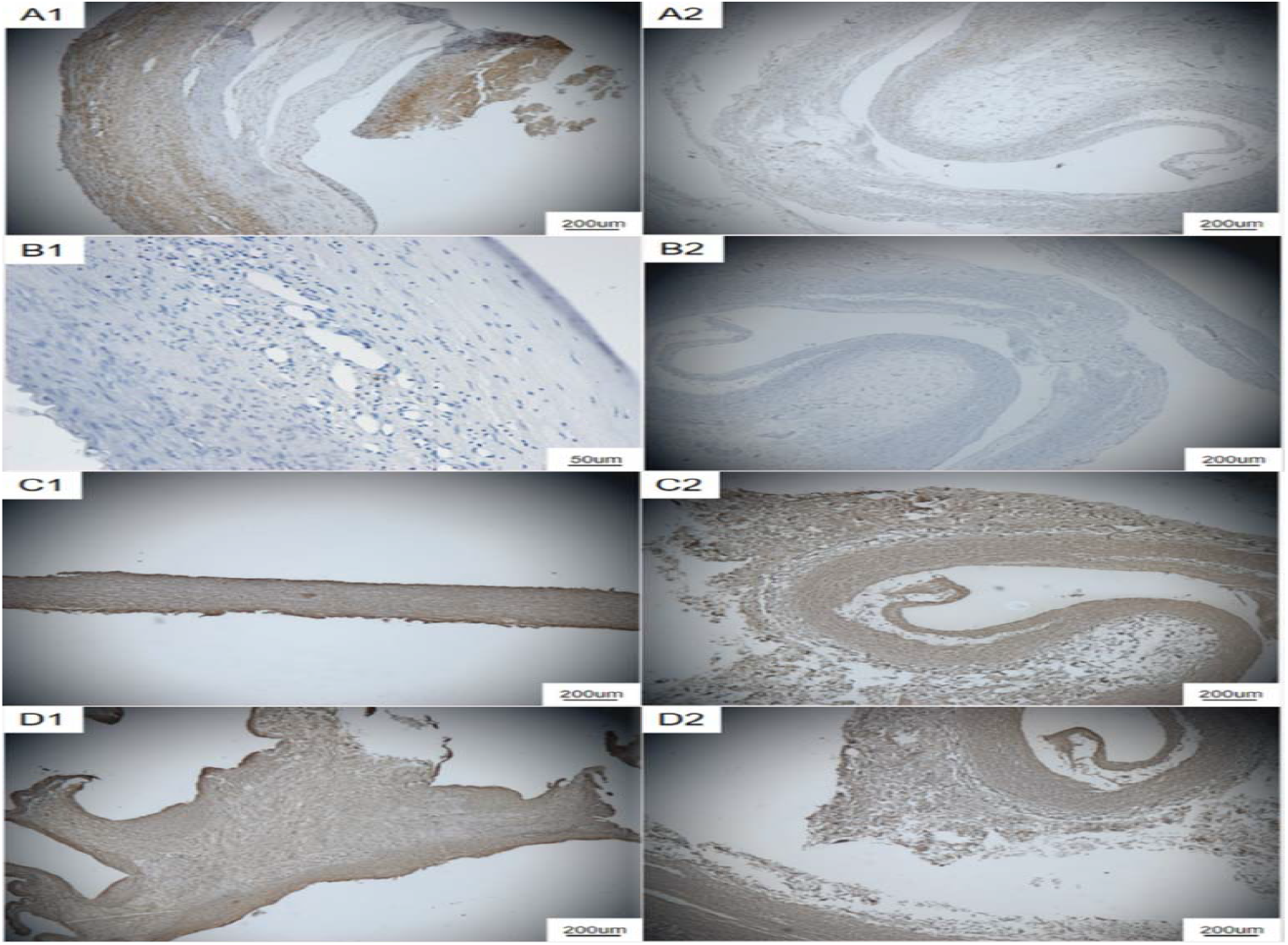
Pathological Staining of Fibrosis-Related Factors. Note: A1: IL-13 staining of pulmonary arterial intimal stripping tissue in CTEPH patients (40x); A2: IL-13 staining of control group pulmonary artery tissue (40x); B1: IL-4 immunohistochemical staining of pulmonary arterial intimal stripping tissue in CTEPH patients (200x); B2: IL-4 staining of control group pulmonary artery tissue (40x); C1: TGF-β1 staining of pulmonary arterial intimal stripping tissue in CTEPH patients (40x); C2: TGF-β1 staining of control group pulmonary artery tissue (40x); D1: SAP immunohistochemical staining of pulmonary arterial intimal stripping tissue in CTEPH patients (40x); D2: SAP staining of control group pulmonary artery tissue (40x).

### Results of Animal Models Simulating Human Conditions

#### Construction of Distal Pulmonary Artery Remodeling Model

By continuously injecting small thrombi into rats on a daily basis, we successfully established a CTEPD rat model of distal pulmonary artery vascular remodeling. HE staining showed thickening of the pulmonary artery and pulmonary microvascular intima in CTEPD rats, with visible reticular and plexiform remodeling (Figure 9). By performing splenectomy and jugular vein cannulation, we successfully established a CTEPD rat model of mid-segment pulmonary artery remodeling (Figure 10). Compared to CTEPD rats without splenectomy, HE staining showed significant remodeling of the mid and distal pulmonary arteries in splenectomized CTEPD rats, with thickened intima, clear boundaries between the intima and media, and narrowed vessel lumens (Figure 11).

**Figure 9.**
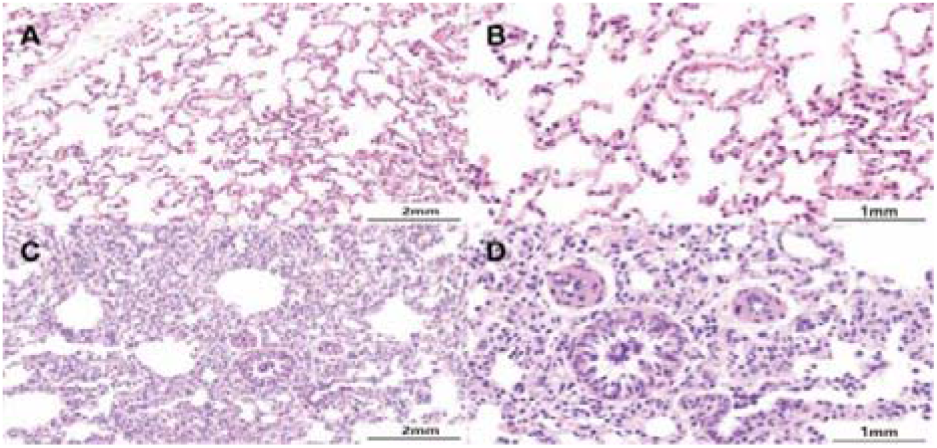
Histopathological HE Staining of Rat Pulmonary Artery Tissue in Different Groups. Note: Sham surgery group 14D (A at 20X magnification, B at 40X magnification) shows no significant changes in the intima. CTEPD group 14D (C at 20X magnification, D at 40X magnification) exhibits obvious thickening of the vessel wall and narrowing of the lumen.

**Figure 10.**
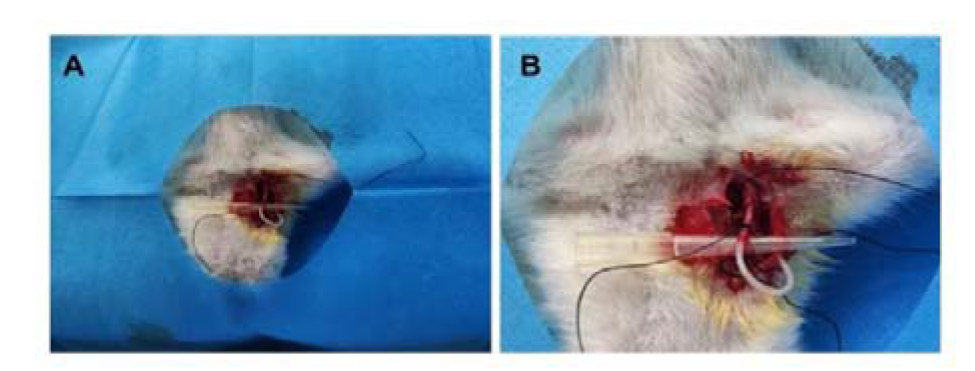
Construction of the Chronic Thromboembolic Pulmonary Hypertension (CTEPH) Rat Model with Central Venous Catheter Placement. Note: A: Ligating the distal end of the left external jugular vein in rats and inserting a central venous catheter; B: Magnified view of the procedure described in A.

**Figure 11.**
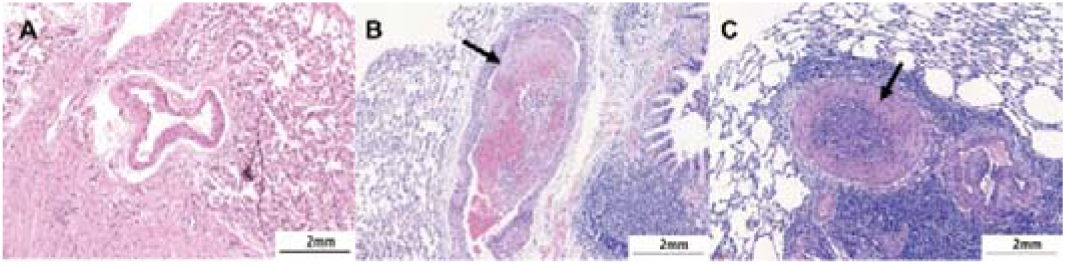
Histopathological HE Staining of Rat Pulmonary Artery Tissue in Different Groups (20X) Note: Sham surgery group 14D (A) shows no significant changes in the intima. CTEPD group 14D without splenectomy (B) exhibits thickening of the vessel wall and slight narrowing of the lumen. CTEPD group 14D with splenectomy (A) shows obvious thickening of the vessel wall and narrowing of the lumen. The black arrows indicate vascular remodeling.

### Construction of Proximal Pulmonary Artery Intimal Fibrosis Model

Using large animals, we successfully established a CTEPD model of intimal fibrosis remodeling by selectively injecting large thrombi into the proximal pulmonary artery. Gross observation of the thrombi in the two lobes of the pulmonary artery after 2 weeks revealed intimal organization, simulating intimal fibrosis caused by proximal vessel occlusion (Figure 12). PTAH staining of the thrombi revealed irregularities in the pulmonary artery lumen, with organized thrombi adhering to the vessel wall and multiple re-canalizations within the thrombi. Proliferative tissue surrounding the pulmonary artery walls encapsulated and divided the thrombi containing blue-purple fibrous protein aggregates.

**Figure 12.**
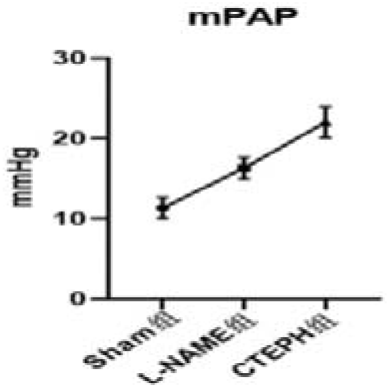
Pressure Curves of Mean Pulmonary Arterial Pressure (mPAP, mmHg) in Different Groups. Note: Sham group: Sham surgery group.

In addition, we successfully established a rat model of proximal pulmonary artery intimal fibrosis in CTEPH rats by intraperitoneal injection of L-NAME and thrombus injection. Measurement of pulmonary artery pressure in each group of rats revealed significantly higher pulmonary artery pressure in CTEPH rats compared to the sham surgery group (P<0.05) and L-NAME group (P<0.05) (see Figure 13, Figure 14, and Table 2). As shown in Figure 15, HE staining showed thickening of the pulmonary artery intima in CTEPH rats, with clear boundaries between the intima and media. Short spindle-shaped fibroblasts were observed in the thickened intima, with small cell heterogeneity, no obvious nucleoli, homogeneous red staining, and a wavy appearance. Sirius red staining and immunohistochemical staining for α-SMA and Elastin revealed the deposition of collagen and other extracellular matrix components in the pulmonary artery intima of CTEPD rats, indicating significant fibrosis (Figure 16).

**Figure 13.**
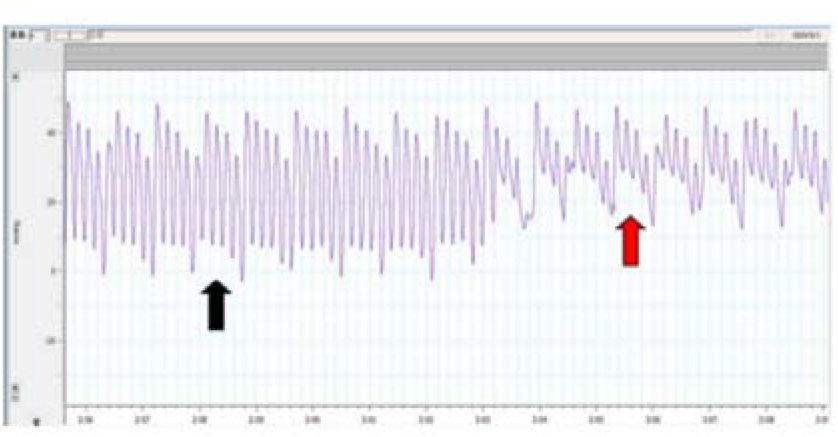
Pressure Curves of Pulmonary Arterial Pressure in Rats Measured by Right Heart Catheterization.

**Figure 14.**
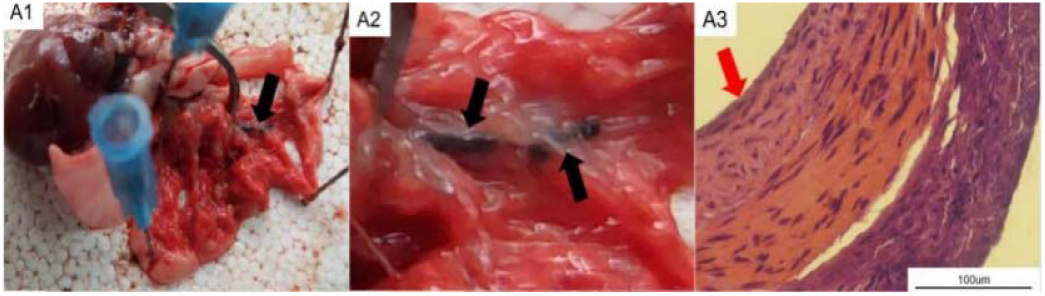
Gross and HE Staining Images of Rat Pulmonary Artery Tissue. Note: A1-A2: Gross appearance of pulmonary tissue in CTEPD rats, with black arrows indicating pulmonary arteries occluded by thrombi. A3: Histopathological HE staining image of pulmonary artery tissue in CTEPD rats, magnification factor is 400x, with red arrows indicating thickened pulmonary artery intima.

**Figure 15.**
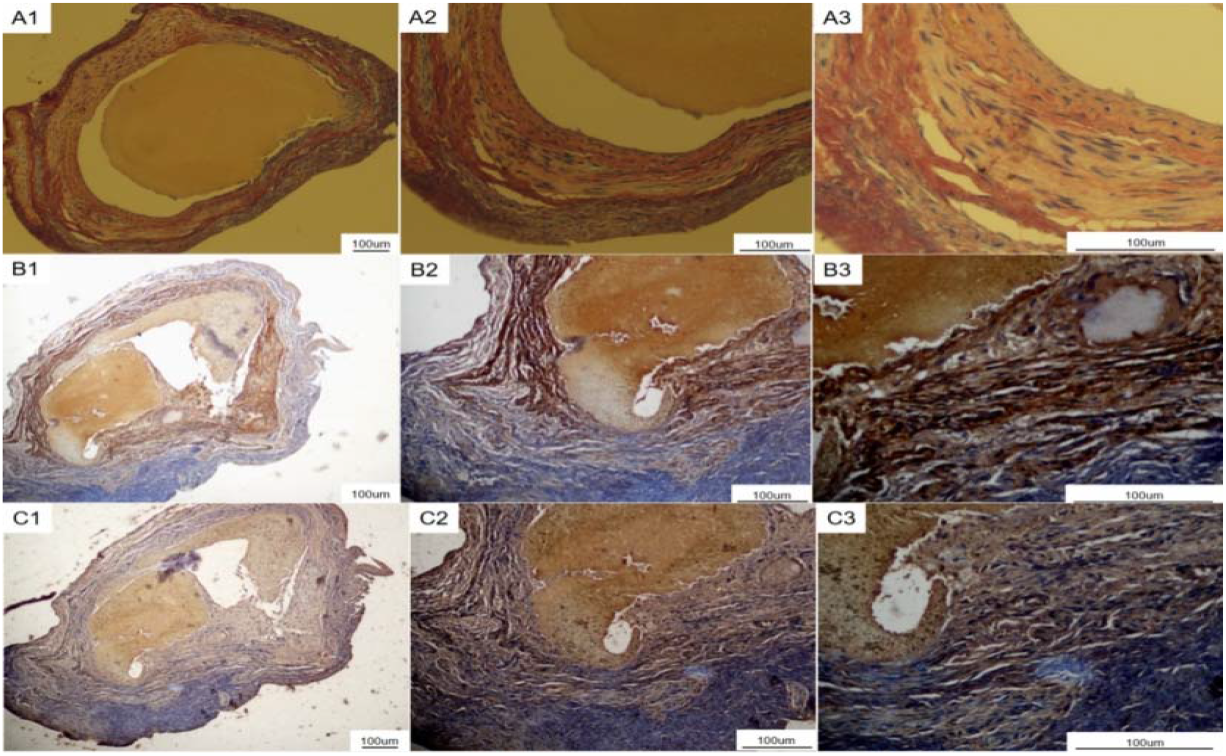
Histopathological Staining Images for Fibrosis Detection in Rat Pulmonary Artery Tissue. Note: A1-A3: Sirius Red staining of pulmonary artery tissue in CTEPD rats, magnification factors are 40x, 100x, and 400x, respectively; B1-B3: α-SMA immunohistochemical staining of pulmonary artery tissue in CTEPD rats, magnification factors are 40x, 100x, and 400x, respectively; C1-C3: Elastin immunohistochemical staining of pulmonary artery tissue in CTEPD rats, magnification factors are 40x, 100x, and 400x, respectively.

**Figure 16.**
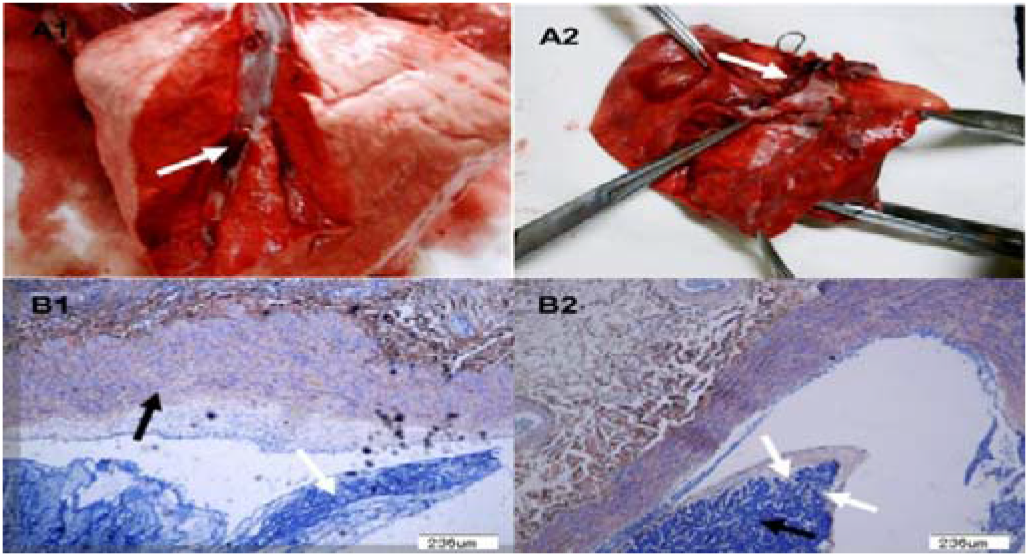
Gross and PTAH Staining Images of Canine Pulmonary Artery Tissue. Note: Image A1: Gross appearance of pulmonary tissue in the 1-week embolism group, white arrows indicate thrombi within the blood vessels. Image A2: Gross appearance of pulmonary tissue in the 2-week embolism group, white arrows indicate intimal organization. Image B1: Thrombi in the 1-week embolism group showing blue-purple stained fibrin polymer (indicated by white arrows) and thickening of the vascular intima (indicated by black arrows), magnification factor is 40x. Image B2: In the 2-week embolism group, proliferation of pulmonary artery wall tissue (indicated by white arrows) is observed, surrounding and dividing the thrombus containing blue-purple stained fibrin (indicated by black arrows), magnification factor is 40x.

**Table 2.**
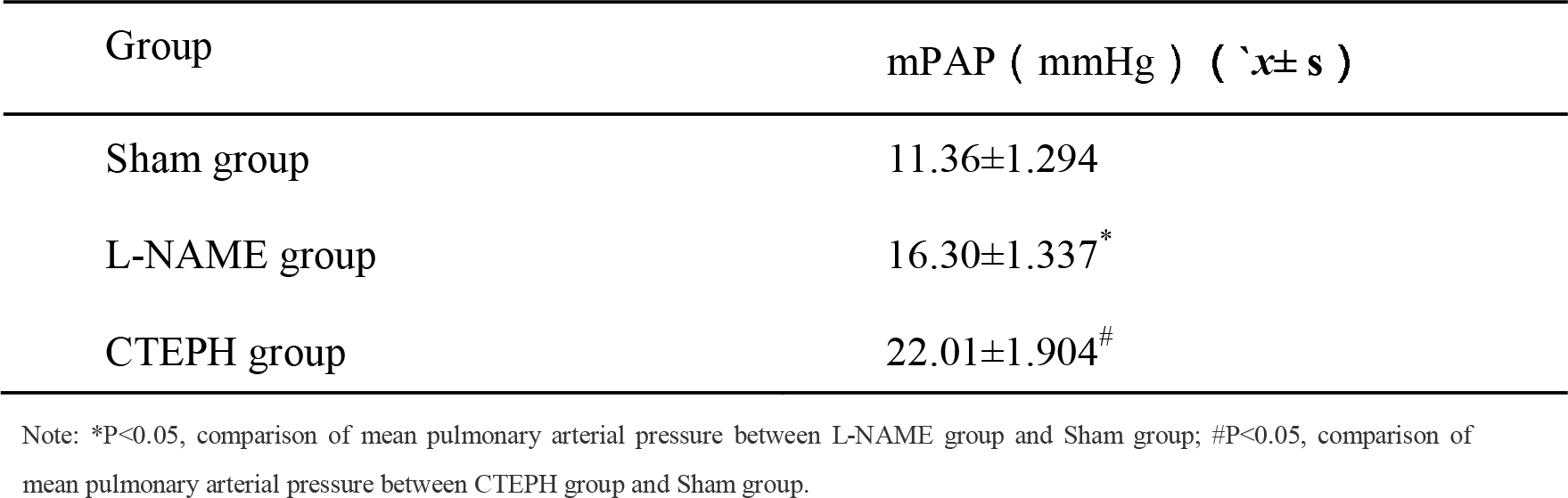
Comparison of mPAP (mmHg) among Different Groups (x± s)

### Construction of Rat Inferior Vena Cava Ligation Model

We constructed an animal model of deep vein thrombosis (DVT) using the inferior vena cava ligation method (Figure 17). HE staining showed residual undissolved thrombi in the clot, with proliferation of short spindle-shaped fibroblasts and small cell heterogeneity, without obvious nucleoli (Figure 18). Masson staining showed a significant positive reaction, indicating increased blue collagen fibers and chronic fibrotic changes in the thrombus (Figure 19).

**Figure 17.**
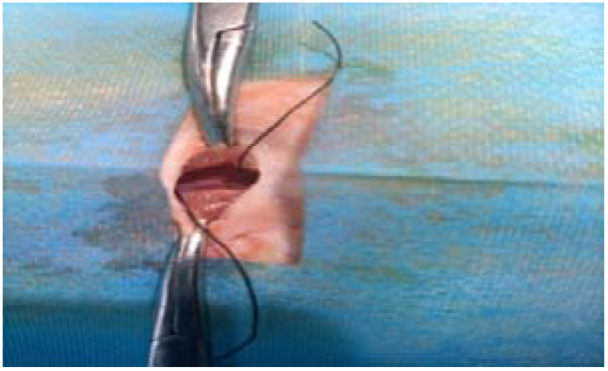
Macroscopic Image of the Surgical Procedure for Establishing a Chronicized Inferior Vena Cava (IVC) Ligation Model in Rats with Deep Vein Thrombosis (DVT).

**Figure 18.**
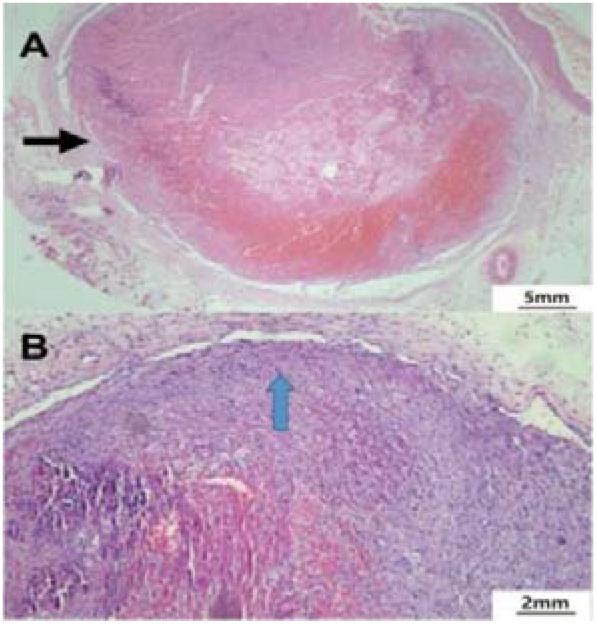
Histopathological HE Staining of Thrombus Tissue in Rat Inferior Vena Cava (IVC) Ligation Model. Note: Image A: HE staining image of thrombus in rat IVC, magnification factor is 4x, black arrows indicate the IVC thrombus. Image B: HE staining image of thrombus in rat IVC, magnification factor is 10x, blue arrows indicate the presence of short spindle-shaped fibroblasts.

**Figure 19.**
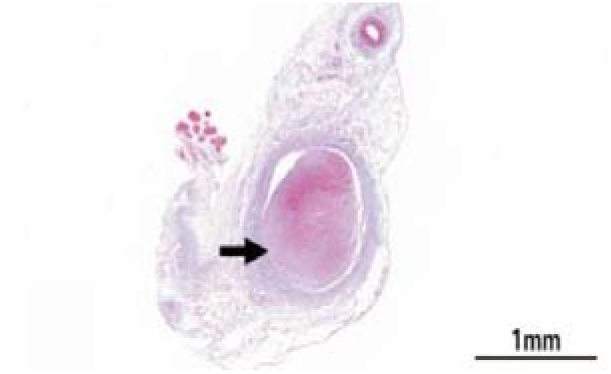
Masson’s Staining of Thrombus Tissue in Rat Inferior Vena Cava (IVC) Ligation Model. Note: The image is magnified at a factor of 40x. Black arrows indicate the IVC thrombus.

### Rats plasma NO levels

The NO content in the normal group of rats was (3.54±0.54) umol/L, while in the chronic pulmonary embolism group of rats, it was (0.15±0.11) umol/L. The difference in NO levels between the two groups was statistically significant (t’ = 15.058, P < 0.001) (Figure 20).

**Figure 20.**
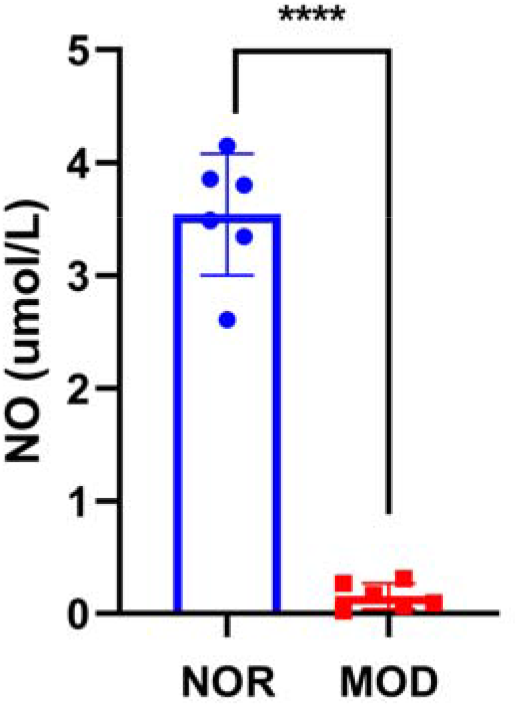
Rats plasma NO levels. Note: NOR:controls.MOD:chronic pulmonary embolism groups. *P<0.05; **P<0.01; ***P<0.001.

## Discussion

CTEPD has a relatively high incidence rate and is difficult to treat due to a lack of understanding of its pathogenesis. Since surgical specimens of PEA represent the end-stage of the disease and it is not possible to dynamically collect tissue samples of DVT-APTE-CTEPD/CTEPH to explore the dynamic evolution of pulmonary vascular lesions, the underlying mechanisms are still unclear. Reliable animal models are powerful tools to explore this process, addressing the current research pain points and difficulties in model construction.

The progression from DVT to APTE-CTEPD/CTEPH is a dynamic process. Under the stimulation of incompletely dissolved thrombi, the intima of the proximal pulmonary artery thickens and fibrosis occurs. Downstream, secondary vascular remodeling of the mid and distal pulmonary arteries occurs due to shear forces and abnormal cytokine secretion, resulting in increased resistance in the pulmonary vascular bed and elevated pulmonary artery pressure [9-11]. In PEA tissues from CTEPD/CTEPH patients, thrombosis organization, recanalization, intimal fibrosis in the proximal pulmonary artery, and vascular remodeling in the mid and distal pulmonary arteries have been observed. There are significant differences in the pathological manifestations among the three regions, and the underlying causes are also different. Therefore, constructing an animal model that can simultaneously simulate the pathological manifestations of the three regions is extremely challenging, and successful models have not been identified [12-16]. The main difficulty lies in the strong fibrinolytic capacity of animals, which often leads to rapid dissolution of injected thrombi, preventing pulmonary vascular remodeling. On the other hand, multiple thrombus injections result in very low animal survival rates. Striking a balance between these two challenges is difficult for researchers. Some researchers have used non-dissolvable polystyrene microspheres injected into the pulmonary artery to induce pulmonary vascular remodeling [17], while others have combined it with left pulmonary artery ligation [18]. Although this does simulate increased pulmonary artery pressure and intimal remodeling, the differences between non-dissolvable microspheres and thrombi raise concerns about the consistency of the intimal remodeling process and the cellular components involved, and it may not adequately explain clinical issues regarding cellular components and molecular mechanisms. Taking into account the factors mentioned above, our strategy is to construct models that simulate different locations and pathological manifestations during the disease process through different modeling approaches, enabling researchers to choose appropriate models based on their specific research directions.

Mid and distal pulmonary arteriole remodeling model: PEA is an effective treatment for CTEPD, but it cannot cure patients with concurrent remodeling of the distal pulmonary arteries. This subgroup of patients has the worst prognosis and poses a challenge in current treatment [19]. Distal pulmonary arteriole remodeling may occur due to two reasons. Firstly, fragmented small thrombi directly obstruct the arterioles and undergo organization, leading to secondary remodeling. Secondly, proximal pulmonary artery obstruction causes redistribution of blood flow and changes in neurohumoral factors and shear forces, resulting in remodeling of the pulmonary arterioles [20]. The latter can be simulated in the proximal pulmonary artery remodeling model, so our distal pulmonary arteriole remodeling model primarily focuses on simulating remodeling caused by fragmented small thrombi. Due to the strong fibrinolytic capacity of animals, small thrombi are usually swiftly dissolved. Therefore, delaying the dissolution of small thrombi is crucial for constructing the distal pulmonary arteriole remodeling model. We know that one of the important functions of the spleen is to filter and remove senescent cells, including red blood cells and platelets, to maintain normal blood viscosity [21]. Studies have shown that a significant number of CTEPD patients have a history of splenectomy, which may be an important risk factor for CTEPD development [22]. Therefore, we attempted to inject small thrombi in combination with splenectomy and successfully delayed the thrombus dissolution in the distal pulmonary arterioles. We observed corresponding remodeling in the mid and distal vessels of the pulmonary artery, including intimal thickening and luminal narrowing. This model not only effectively simulates the pathological manifestations of distal pulmonary arteriole remodeling but also mimics the dynamic changes in chronic pulmonary embolism in patients with splenectomy. Overall, this model is suitable for studying the dynamic changes, cellular components, and molecular mechanisms involved in distal pulmonary arteriole remodeling during the chronicization process of pulmonary embolism.

Proximal pulmonary artery intimal fibrosis model: Thrombus organization and subsequent intimal remodeling in the proximal pulmonary arteries are the triggering points and main causes of CTEPD development. This process is central to the chronicization of pulmonary embolism, secondary remodeling of the pulmonary vascular bed, and increased pulmonary artery pressure. It is also the primary focus and necessary path for mechanistic research. Our study found a significant deposition of extracellular matrix and pronounced fibrosis in the proximal pulmonary artery intima. This extracellular matrix is mainly derived from a large number of α-SMA+ myofibroblasts present in the intima. These findings are consistent with previous studies, but the origin of myofibroblasts remains unclear [23-25]. In order to explore the possible source of α-SMA+ myofibroblasts, we performed immunofluorescent double staining and identified a large number of fibroblasts in the intima of CTEPH patients’ proximal pulmonary arteries. Their transformation may be related to the abundant expression of IL-13 in the intima. However, even so, we still lack a clear understanding of the specific transformation mechanism and dynamic evolutionary process.

Resolving these questions relies on a mature animal model that allows for dynamic observation of pulmonary artery remodeling. In order to simulate the remodeling of the proximal pulmonary artery, it is necessary to trap the thrombus in a large pulmonary artery and prevent it from being washed away by blood flow. However, the pulmonary artery has a larger lumen and is significantly more extensible than typical arteries. Therefore, the thrombus blocking the main trunk of the proximal pulmonary artery must be large enough and remain in place for a sufficient duration, with the latter being the main challenge. We simulated the chronicization process of thrombus in the proximal pulmonary artery by selectively occluding the proximal pulmonary artery in dogs [26]. The manifestations included thrombus organization, cellular ingrowth into the thrombus, segmentation, and encasement of the thrombus. These are generally considered as early signs of recanalization, and the intima of the pulmonary artery markedly thickened, resembling the peeled-off intimal tissue in CTEPH patients. This model effectively represents the pathological changes of chronicization in the proximal pulmonary artery thrombus. However, its drawbacks include the high cost of experimental dogs, low pulmonary artery pressure, and the technical difficulty of selective occlusion. We recommend that laboratories with the necessary conditions consider using this approach for modeling. Thus, we attempted to establish a more cost-effective rat model. We selected SD rats, which have lower costs and less demanding housing requirements, as the subjects for our study. However, the strong fibrinolytic capacity of rats often leads to rapid dissolution of injected thrombi, so delaying their dissolution is crucial for modeling.

We started by investigating the mechanism of thrombus dissolution and found that nitric oxide (NO) plays an important role in the process. Previous studies have shown that endogenous nitric oxide synthase inhibitors in the peripheral blood of CTEPH patients are significantly increased compared to healthy controls [27]. After thrombus occlusion, NO is produced by endothelial cells, mediating vasodilation and promoting thrombus dissolution [28]. Therefore, we attempted to inhibit the production of NO by endothelial cells using a nitric oxide synthase inhibitor (NOSI) to block the thrombus-dissolving effect of NO. The results showed that the NOSI group in the rat model had a significantly prolonged thrombus retention time compared to the control group without NOSI. The large thrombus in the occluded pulmonary artery underwent organization, fibrotic transformation, and significant intimal thickening. Sirius Red staining confirmed marked intimal fibrosis in the pulmonary artery, and we even observed cellular ingrowth into the thrombus, which may be a process of vascular recanalization that requires further exploration in subsequent studies. Additionally, the animal model constructed using this approach exhibited significantly increased pulmonary artery pressure compared to normal rats. In summary, the model we constructed effectively simulates the process of intimal fibrosis in the pulmonary artery caused by the chronicization of thrombus in patients with acute pulmonary embolism. Its pathological manifestations resemble the intima of CTEPH patients, and it simulates the pathophysiological features of CTEPH. This model has the potential to become a powerful tool for exploring the mechanisms underlying pulmonary artery remodeling during the evolution from acute pulmonary thromboembolism to the chronicization of pulmonary embolism.

Inferior vena cava ligation model: The source of progression from DVT-APTE-CTEPD/CTEPH in clinical practice is deep venous thrombosis that detaches and evolves into CTEPH [29]. Therefore, our model for studying the chronicization of CTEPH thrombus is established by ligating the inferior vena cava in rats to induce in situ thrombus formation. The biggest advantage of this model compared to the previous one is that it is the only model that does not require ex vivo manipulation of the thrombus. The cellular components within the thrombus remain consistent with the pulmonary artery thrombus cellular components in patients with chronic pulmonary embolism [30]. We can refer to this as a "live thrombus," whereas the thrombus used in the previous model undergoes ex vivo manipulation, leading to cell death and changes in composition, which can be referred to as a "dead thrombus." This model does not require the injection of artificial thrombus, thereby eliminating interference caused by changes in cellular components due to ex vivo manipulation of the thrombus. Using this model, we can most accurately reflect the dynamic evolution process of thrombus within the vessel lumen, track the cellular components, and investigate the molecular mechanisms involved in its evolution. This model presents a more complete representation of the thrombus evolution process, from fresh thrombus formation to thrombus fibrosis and subsequent dissolution and recanalization. Therefore, it provides a basis for studying the cellular components and molecular mechanisms of thrombus organization and recanalization processes.

These different animal models provide various aspects for studying the pathophysiological mechanisms of CTEPH. Researchers can choose the appropriate model from the above options based on their specific research purposes, directions, and requirements.

## Conclusion, Limitations, and Outlook

In summary, the animal models of chronic progression of DVT-APTE-CTEPD/CTEPH that we constructed provide a pathological and physiological basis for understanding the remodeling of the pulmonary artery, distal vascular changes, pulmonary artery intimal fibrosis, and chronicization of pulmonary embolism thrombus in the progression of VTE. These models demonstrate pathological manifestations that are consistent with the chronicization of thrombus observed in humans. However, currently, these animal models can only simulate the chronicization of VTE thrombus and do not accurately replicate the significant increase in pulmonary artery pressure observed in CTEPH. Therefore, establishing a stable CTEPH animal research model still requires the combination of scientifically effective factors such as the infusion of larger autologous thrombus through external jugular vein cannulation, administration of L-NAME, and splenectomy. However, it is necessary to exclude the additional influence of these multiple factors on the progression of VTE. Only then can we explore a VTE model that is more clinically relevant and establish a stable animal model that mimics the pathological and physiological changes observed in humans for studying intimal stripping in CTEPH in the future.

## Authors’ contributions

All authors participated in the design and interpretation of the studies, the analysis of the data, and the review of the manuscript. CS Deng made a substantial contribution to the conception of the study, conducted the experiments, and acquired data; QH Lin, WF Wang and XY Chen revised the paper for intellectual content; N Shao, QX Wu and XY Lai helped establish the animal model and prepare the autologous blood clots; MH Chen, M Chen and YJ Wu recorded the parameters and stained the specimens; DW Wu and HL Li performed the statistical analysis. All authors gave final approval for publication.

